# Mutation Y453F in the spike protein of SARS-CoV-2 enhances interaction with the mink ACE2 receptor for host adaption

**DOI:** 10.1101/2021.08.24.457448

**Authors:** Wenlin Ren, Jun Lan, Xiaohui Ju, Mingli Gong, Quanxin Long, Zihui Zhu, Yanying Yu, Jianping Wu, Jin Zhong, Rong Zhang, Shilong Fan, Guocai Zhong, Ailong Huang, Xinquan Wang, Qiang Ding

## Abstract

COVID-19 patients transmitted SARS-CoV-2 to minks in the Netherlands in April 2020. Subsequently, the mink-associated virus (miSARS-CoV-2) spilled back over into humans. Genetic sequences of the miSARS-CoV-2 identified a new genetic variant known as “Cluster 5” that contained mutations in the spike protein. However, the functional properties of these “Cluster 5” mutations have not been well established. In this study, we found that the Y453F mutation located in the RBD domain of miSARS-CoV-2 is an adaptive mutation that enhances binding to mink ACE2 and other orthologs of *Mustela* species without compromising, and even enhancing, its ability to utilize human ACE2 as a receptor for entry. Structural analysis suggested that despite the similarity in the overall binding mode of SARS-CoV-2 RBD to human and mink ACE2, Y34 of mink ACE2 was better suited to interact with a Phe rather than a Tyr at position 453 of the viral RBD due to less steric clash and tighter hydrophobic-driven interaction. Additionally, the Y453F spike exhibited resistance to convalescent serum, posing a risk for vaccine development. Thus, our study suggests that since the initial transmission from humans, SARS-CoV-2 evolved to adapt to the mink host, leading to widespread circulation among minks while still retaining its ability to efficiently utilize human ACE2 for entry, thus allowing for transmission of the miSARS-CoV-2 back into humans. These findings underscore the importance of active surveillance of SARS-CoV-2 evolution in *Mustela* species and other susceptible hosts in order to prevent future outbreaks.

## Introduction

Coronaviruses are enveloped, positive-stranded RNA viruses that circulate broadly among humans, other mammals, and birds and can cause respiratory, enteric, and hepatic disease^1^. In the last two decades, coronaviruses have caused three major outbreaks: severe acute respiratory syndrome (SARS), Middle East respiratory syndrome (MERS) and the recent Coronavirus Disease 2019 (COVID-19)^2,3^. COVID-19, caused by severe acute respiratory syndrome coronavirus 2 (SARS-CoV-2), is a major global health threat.

The receptor-binding domain (RBD) of the SARS-CoV-2 spike (S) protein binds its cellular receptor angiotensin-converting enzyme 2 (ACE2), thus mediating viral entry^4,5^. It has been demonstrated that the interaction of a virus with (a) species-specific receptor(s) is a primary determinant of host tropism and constitutes a major interspecies barrier at the level of viral entry^6^. Our previous study found that numerous mammalian ACE2 orthologs could function as receptors to mediate virus entry *in vitro*^7^, and other studies have demonstrated that rhesus macaques, dogs, cats, cattle, hamsters, ferrets, minks and other animals are susceptible hosts^8–13^. Together, these findings suggest that SARS-CoV-2 has a broad host range, with many species that could serve as potential reservoirs and thus pose a risk for spillover to humans in the future^7^.

Recently, the first animal-to-human transmission of SARS-CoV-2 was reported^14,15^. In April 2020, SARS-CoV-2 was transmitted to minks at two farms in the Netherlands by infected employees, and the virus subsequently circulated among the minks^16,17^. On 5 November 2020, the Danish public health authorities reported 214 cases of humans infected with mink-associated SARS-CoV-2 variants (miSARS-CoV-2) containing a combination of mutations not previously observed^15^. Genetic analysis grouped the miSARS-CoV-2 variants into 5 clusters with seven mutations^18^. Of note, the “Cluster 5” variant with four amino acid changes in the spike protein was identified in mink and isolated from 12 human cases in North Jutland^14,18^. The implications of the mutations in this variant are not yet well characterized. Preliminary results suggested that the “Cluster 5” miSARS-CoV-2 strain has moderately decreased sensitivity to neutralizing antibodies^14,19,20^. Further investigations are urgently needed to dissect the biological significance of these mutations and to understand the implications of these identified changes for diagnostics, therapeutics and vaccine development.

In this study, we demonstrate that the Y453F mutation in the miSARS-CoV-2 spike is an adaptive mutation that increases interaction with mink ACE2 without compromising utilization of human ACE2. In addition, the miSARS-CoV-2 exhibited partial resistance to neutralization with convalescent serum. Our study not only provides critical insights into viral adaption and evolution but also highlights the importance of surveillance of viral variants in animals to reduce the risk of the emergence of new viral variants and prevent future outbreaks.

## RESULTS

### Potential impact of the Y453F substitution on the binding of SARS-CoV-2 spike to mink ACE2

Sequence analysis of the miSARS-CoV-2 “Cluster 5” variant isolated from patients identified four mutations in the spike protein: del 69–70 (a deletion of the His69 and Val70), Y453F (located in the RBD), I692V, and M1229I^14^ (**Fig. 1A**). Based on the structure of SARS-CoV-2 spike bound to human ACE2^21–24^, Y453 in the spike protein can form hydrogen bonds with H34 in human ACE2 (**Fig. 1B and C, left panel**); however, mink ACE2 has a Tyr at the 34th position, which is predicted to cause a spatial clash with Y453 (**Fig. 1B and C, right panel**) and impair interaction with mink ACE2. This is supported by the limited binding affinity observed between WT SARS-CoV-2 RBD and mink ACE2 (**Fig. 2A and B**), and the finding that mink ACE2 mediates authentic SARS-CoV-2 entry with 20-30% efficiency of human ACE2^7^. Based on this analysis, we hypothesized that the Y453F substitution evolved in the miSARS-CoV-2 spike RBD as it enhances interaction with mink ACE2, conferring a fitness advantage in the new host and leading to circulation of the adapted virus in the mink population.

**Figure 1.**
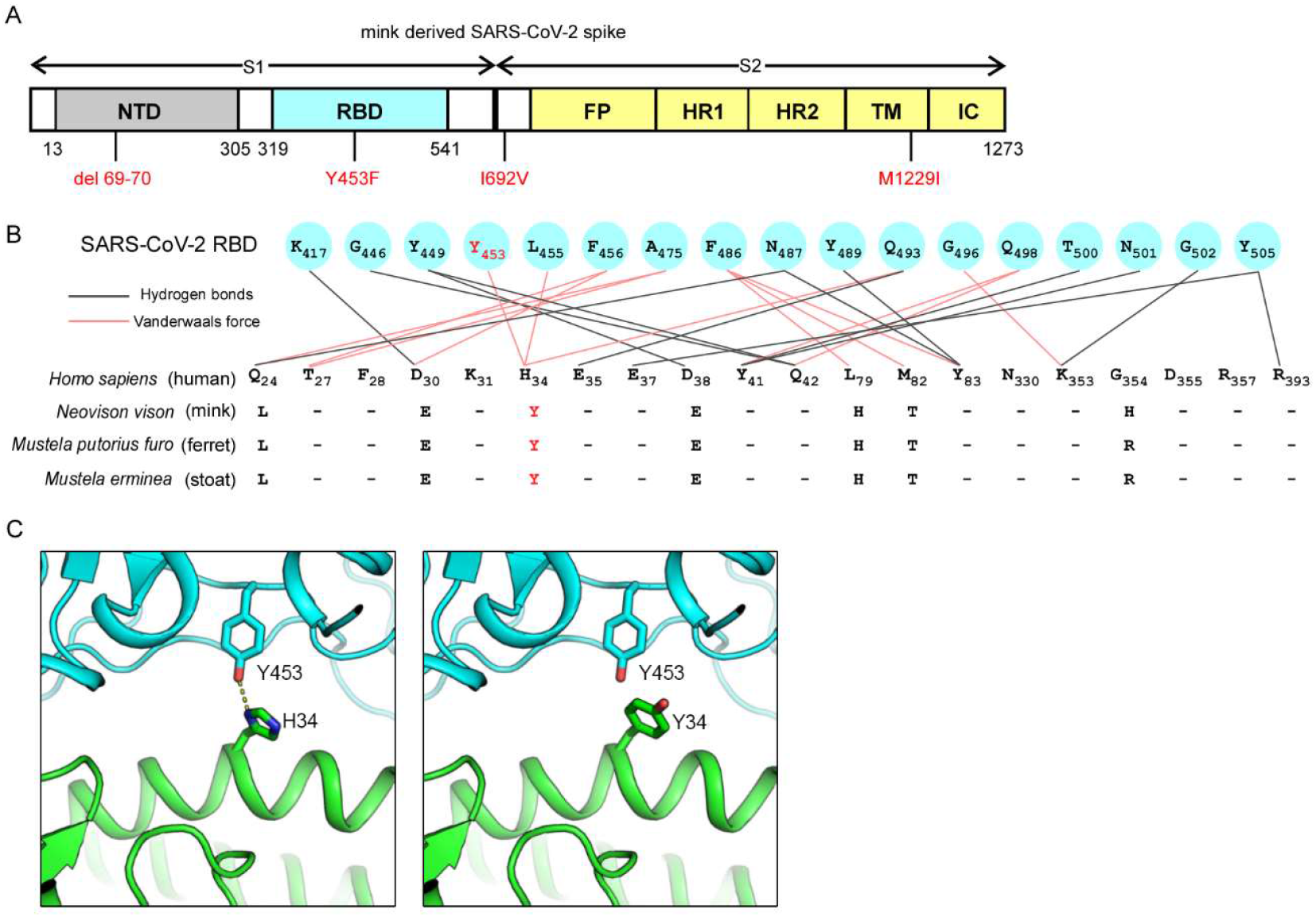
The mutation Y453F in SARS-CoV-2 spike protein is a potential genetic adaptation in minks. (A) Schematic of amino acid changes in the spike protein of miSARS-CoV-2 variant “Cluster 5”. The four genetic changes in the S protein are marked in red. NTD, N-terminal domain; FP, fusion peptide; HR1, heptad repeat 1; HR2, heptad repeat 2; TM, transmembrane region; IC, intracellular domain. (B) Alignment of the residues of human (*Homo sapiens*; NCBI Reference Sequence: NM_001371415.1), mink (*Neovison vison*; NCBI Reference Sequence: MW269526.1), ferret (*Mustela putorius furo*; NCBI Reference Sequence: NM_001310190.1) and stoat (*Mustela erminea*; NCBI Reference Sequence: XM_032331786.1) ACE2 at the interface of ACE2 with the SARS-CoV-2 spike protein. The position of Y453 in the SARS-CoV-2 RBD is colored in red. The Y34 of mink, ferret and stoat ACE2 are highlighted in red. (C) Structural features of the SARS-CoV-2 RBD and ACE2. Contacting residues are shown as sticks at the hACE2-SARS-CoV-2 RBD interfaces (PDB ID: 6M0J). The key residues Y453 of the RBD and H34 of human ACE2 (left) are labeled. The position of the Y34 substitution in human ACE2 (right) is labeled. SARS-CoV-2 RBD is shown in cyan, ACE2 in green. The dashed lines indicate hydrogen bonds.

**Figure 2.**
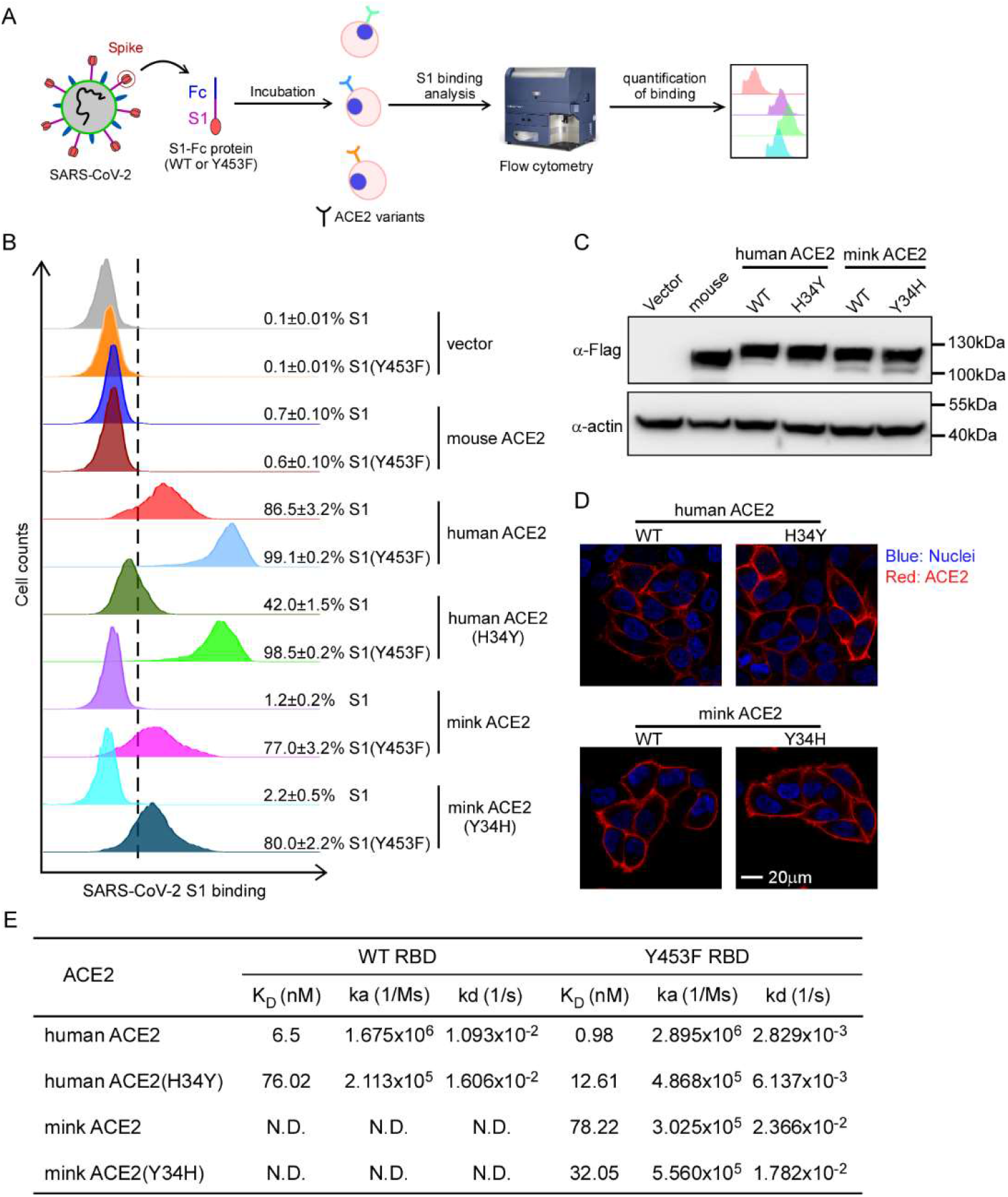
Increased binding of Y453F spike protein to mink ACE2. (A) Schematic of testing the efficiency of ACE2 variants binding WT or Y453F viral spikes. (B) HeLa cells were transduced with human and mink ACE2 or their mutants as indicated. The transduced cells were incubated with the WT or Y453F S1 domain of SARS-CoV-2 C-terminally fused with a His tag and then stained anti-His-PE for flow cytometry analysis. Values are expressed as the percent of cells positive for S1-Fc among the ACE2-expressing cells (zsGreen1+ cells) and shown as the means ± SD from 3 biological replicates. These experiments were independently performed three times with similar results. (C) Representative immunoblots of HeLa cells transduced with lentiviruses expressing FLAG-tagged ACE2. Actin used as the loading control. These experiments were independently performed twice with similar results. (D) Cell surface localization of human and mink ACE2 and their mutants. HeLa cells were transduced with lentivirus expressing human or mink ACE2 and their mutants. Cells were incubated with rabbit polyclonal antibody against ACE2 and then stained with goat anti-rabbit IgG (H+L) conjugated with Alexa Fluor 568 and DAPI (1μg/ml). The cell images were captured with a Zeiss LSM 880 Confocal Microscope. This experiment was independently repeated twice with similar results and the representative images are shown. (E) Table summarizing biochemical results for ACE2 variants bound to WT or Y453F RBD. These experiments were independently performed three times with similar results.

### The Y453F mutation in miSARS-CoV-2 spike increases interaction with mink ACE2

To test our hypothesis, we first employed a cell-based assay, using flow cytometry to assess the binding of S protein to ACE2 orthologs and variants (**Fig. 2A**). We cloned the cDNA of ACE2 variants, each with a C-terminal FLAG tag, into a bicistronic lentiviral vector that expresses the fluorescent protein zsGreen1 via an IRES element (pLVX-IRES-zsGreen1) and can be used to monitor transduction efficiency. Next, WT or Y453F S1-Fc (a purified fusion protein consisting of the S1 domain of SARS-CoV-2 S protein and an Fc domain of human lgG) was incubated with HeLa cells transduced with the ACE2 variants. Binding of S1-Fc to ACE2 was then quantified by flow cytometry as the percent of ACE2^-^ expressing cells positive for S1-Fc (**Fig. S2** and **Fig. 2B**).

As expected, the binding of S1-Fc or S1 (Y453F)-Fc to HeLa cells expressing mouse ACE2 was very low and comparable to that of the empty vector control while S1-Fc efficiently bound to HeLa cells expressing human ACE2, which is consistent with previous reports^5^. S1 (Y453F)-Fc bound human ACE2 more efficiently than WT S1-Fc (99.1% vs 86.5%). Notably, WT S1-Fc bound mink ACE2 with limited efficiency, but in contrast, S1 (Y453F)-Fc bound mink ACE2 with 77% efficiency, demonstrating that the miSARS-CoV-2 mutation enhances binding to mink ACE2. Moreover, after replacing the amino acid residue at position 34 of human ACE2 with its mink counterpart to generate human ACE2 (H34Y), binding to S1-Fc was reduced (86.5% [WT] vs 42.0%) but increased to S1 (Y453F)-Fc (92.25% [WT] vs 98.38%). Performing the converse by substituting the amino acid residue at position 34 of mink ACE2 with its human counterpart to generate mink ACE2 (Y34H) only slightly increased binding to S1-Fc (0.04% vs 0.92%) and decreased binding to S1(Y453F)-Fc (45.05% vs 42.17%) (**Fig. 2B**).

As we used cell-surface staining of ACE2 to sort the cells used for these experiments so that they had comparable ACE2 expression, the limited or undetectable binding of certain ACE2 variants with the S1 variants was not due to low expression of ACE2 or alteration of its cell surface localization (**Fig. S1A**). The expression level of the ACE2 variants was also assessed by immunoblotting using an anti-Flag antibody and cell surface localization by immunofluorescent microscopy (**Fig. 2C and D**). Together, these results showed that all the ACE2 orthologs were expressed and localized at the cell surface at comparable levels, excluding the possibility that the limited binding efficiencies of ACE2 orthologs with S1-Fc variants was attributable to varied cell surface localization.

To further quantify the binding of ACE2 variants with the spike protein variants, we expressed and purified recombinant WT and Y453F SARS-CoV-2 RBD as well as ACE2 variants to assay binding *in vitro* by surface plasmon resonance (SPR) analysis (**Fig. S3A-B** and **Fig. 2E**). Mink ACE2 bound Y453F SARS-CoV-2 with a K_D_ of 78.22nM but binding to WT RBD was not detectable. Human ACE2 bound SARS-CoV-2 RBD with a K_D_ of 6.5nM, and Y453F RBD with a K_D_ of 0.98nM. These SPR results are thus consistent with the findings of our cell-based assay.

Collectively, our results demonstrate that the SARS-CoV-2 spike binds mink ACE2 with limited affinity. However, the Y453F mutation dramatically increased the binding affinity with mink ACE2 without compromising binding to human ACE2, which suggests that Y453F is an adaptive mutation to improve its fitness in a new host—mink.

### Structural basis for the enhanced binding of mink ACE2 with Y453F RBD

To elucidate at the atomic level the molecular basis for the enhanced binding, we determined the crystal structure of SARS-CoV-2 Y453F RBD bound to mink ACE2 at 3.01 Å resolution (**Table 1**). The overall binding mode of Y453F RBD to mink ACE2 was very similar to that of WT RBD to human ACE2^21^ (**Fig. 3A and Table S1**) as evidenced by the low root mean square deviation (RMSD) value of 0.9 Å for the aligned 597 Cα atoms. A surface of 1668 Å^2^ is buried at the binding interface of Y453F RBD/mink ACE2 and is comprised of 18 residues from Y453 RBD and 18 residues from mink ACE2. The WT RBD/human ACE2 binding interface has a similar buried surface of 1687 Å^2^, made of 17 WT RBD residues and 20 human ACE2 residues.

**Table 1.**
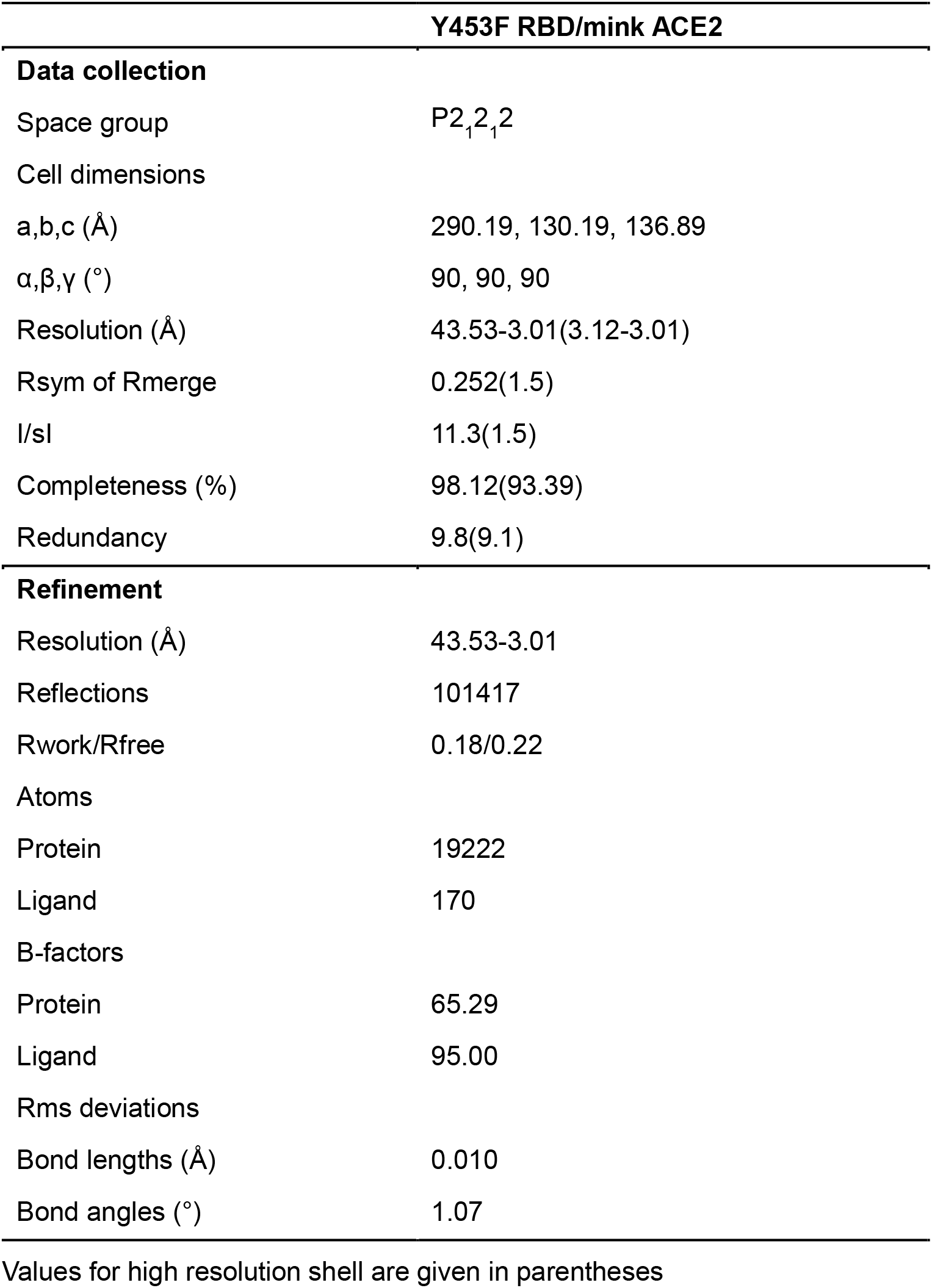
Data collection and refinement statistics.

**Figure 3.**
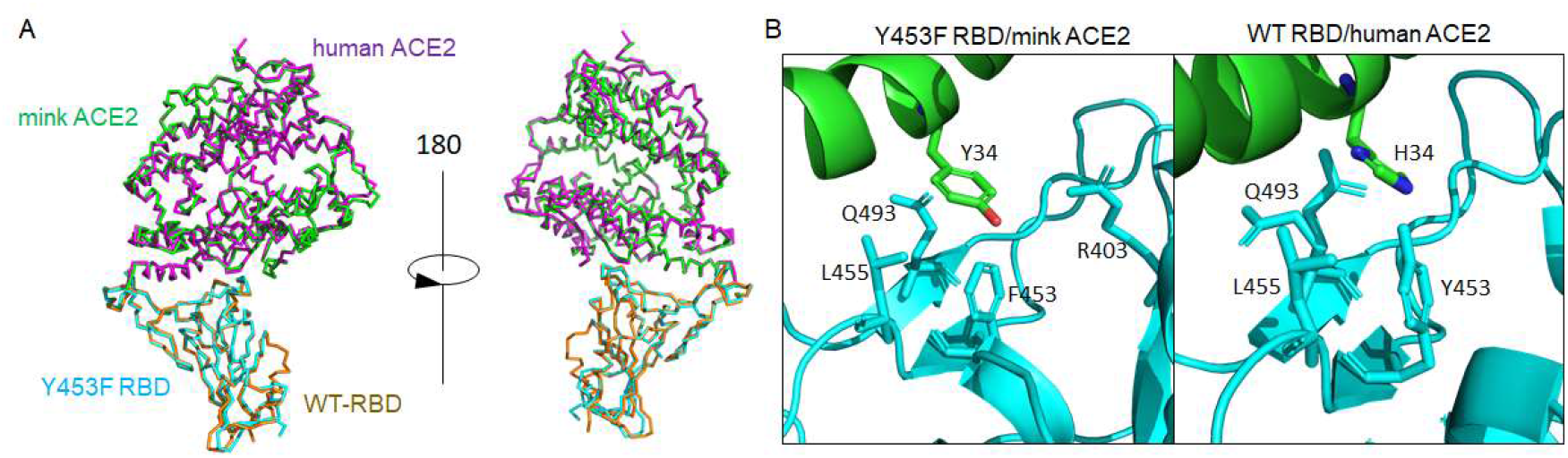
Structural comparison of Y453F RBD/mink ACE2 and WT RBD/human ACE2 complexes. (A)The overall structure is shown as ribbons. The Y453F-RBD is shown in cyan and mink ACE2 in green. The WT-RBD and human ACE2 are shown in magenta and orange, respectively. (B) Interaction residues around position 34 of human and mink ACE2. Left: complex of Y453F RBD/mink ACE2. Right: complex of WT RBD/human ACE2. RBD is shown in cyan, ACE2 in green. Contact residues are shown as sticks which were defined using a distance cutoff of 4Å. Q493 of WT-RBD has a double conformation. The PDB code for WT-RBD/human ACE2 complex is 6M0J^21^.

Between mink ACE2 and human ACE2, there are differences in five of the amino acids involved in RBD binding: L24 (mink ACE2)/Q24 (human ACE2), E30/D30, Y34/H34, E38/D38 and H354/G354 (**Table S1**). We especially focused on the residue at ACE2 position 34 as it directly interacts with RBD residues including that at position 453. Structural analysis indicated that the Y453F substitution in the spike RBD is a species-specific adaptive mutation increasing the binding to mink ACE2 (**Fig. 3B and Table S1**). At the WT RBD/human ACE2 interface, RBD Y453 interacts with H34 of human ACE2 and the change to Tyr, as seen in mink ACE2, results in a bulkier side chain that may increase steric clashes and disfavor the binding of WT RBD to mink ACE2 (**Fig. 3B**). However, with the mutation to Phe at position 453 in the RBD, the Tyr at position 34 in mink ACE2 does not face the same steric clashes (**Fig. 3B**). In addition, while the human ACE2 H34 also has interactions with WT RBD L455 and Q493, mink ACE2 Y34 interacts with residue R403 in addition to L455 and Q493 (**Fig. 3B and Table S1**). These differing interactions may account for the enhanced binding of Y453F RBD to mink ACE2. Finally, it is also predicted that the Y453F mutation would not bring steric clashes with human ACE2 H34 and thus would not significantly change the surrounding interactions, which may explain the retained binding of Y453F RBD to human ACE2.

### The Y453F mutation in miSARS-CoV-2 spike increases its interaction with other *Mustela* ACE2 orthologs

Mink is a member of the *Mustela* genus which also includes stoats and ferrets, the latter of which have been used as animal models owing to their susceptibility to SARS-CoV-2^25^. Due to the high similarity of ACE2 proteins in the *Mustela* genus (**Fig. S4A**), we performed the binding experiments of the S1 variants (WT or Y453F) with ferret and stoat ACE2. Consistent with the results of mink ACE2, ferret and stoat ACE2 exhibited limited binding capability with WT S1-Fc but increased binding ability with S1 (Y453F)-Fc (**Fig. 4A**). As before, we confirmed that the ACE2 orthologs expressed and localized at the cell surface at comparable levels to exclude the possibility that differences in binding capability were due to variation in ACE2 expression and/or mislocalization (**Fig.4B and Fig. S4B**). In line with our cell-based binding assay, SPR analysis did not detect the binding of WT RBD with ferret or stoat ACE2, but Y453F RBD could bind ferret or stoat ACE2 with a K_D_ of 31.17nM and 48.06 nM, respectively (**Fig. 4C**). Taken together, all of these results suggest that the Y453F mutation increases SARS-CoV-2 RBD binding affinity with *Mustela* ACE2 orthologs.

**Figure 4.**
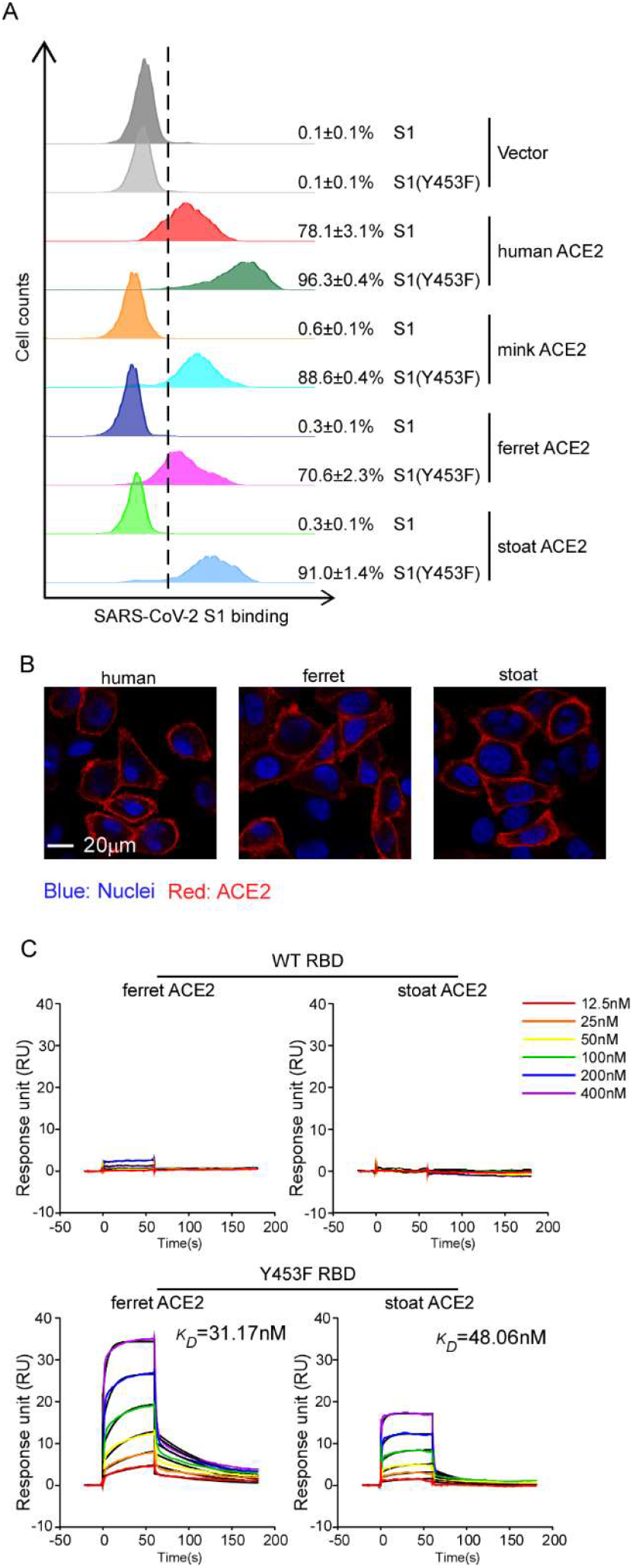
Increased binding of Y453F spike protein to *Mustelidae* ACE2 orthologs. (A) HeLa cells were transduced with human or *Mustelidae* ACE2 orthologs as indicated. Transduced cells were incubated with WT or Y453F S1 domain of SARS-CoV-2 C-terminally fused with His tag and then stained with anti-His-PE for flow cytometry analysis. Values are expressed as the percent of cells positive for S1-Fc among the ACE2-expressing cells (zsGreen1+ cells) and shown as the means ± SD from 3 biological replicates. These experiments were independently performed three times with similar results. (B) Cell surface localization of human, ferret and stoat ACE2. HeLa cells were transduced with lentivirus expressing ACE2 orthologs as indicated. Cells were incubated with rabbit polyclonal antibody against ACE2 and then stained with goat anti-rabbit IgG (H+L) conjugated with Alexa Fluor 568 and DAPI (1μg/ml). The cell images were captured with a Zeiss LSM 880 Confocal Microscope. This experiment was independently repeated twice with similar results and the representative images are shown. (C) The binding kinetics of ACE2 proteins (ferret or stoat) with recombinant WT or Y453F SARS-CoV-2 RBD were obtained using the BIAcore. ACE2 proteins were captured on the chip, and serial dilutions of RBD were then injected over the chip surface. Experiments were performed three times with similar result, and one set of representative data is displayed.

### Y453F spike utilized mink ACE2 with enhanced efficiency for cell entry

To further demonstrate the biological consequences of the enhanced binding affinity of S1 (Y453F) with *Mustela* ACE2, we generated MLV retroviral particles (Fluc as the reporter) pseduotyped with SARS-CoV-2 WT spike (SARS-CoV-2pp) or variant Y453F (SARS-CoV-2pp Y453F) (**Fig. 5A**). A549 cells transduced with ACE2 orthologs were inoculated with the pseudotyped virus, and luciferase (Luc) activity assayed 48h later to determine entry efficiency. Mouse ACE2 was included as a negative control. As expected, Luc activities were limited in A549-mouse ACE2 cells infected with WT or Y453F SARS-CoV-2pp. However, Luc activity increased by 350 and 400 fold in A549-human ACE2 cells infected with WT or Y453F SARS-CoV-2pp, respectively, compared to that of A549-mouse ACE2. In mink or ferret ACE2-transduced cells infected with SARS-CoV-2pp, the Luc activity increased by 9 or 12 fold relative to that of mouse ACE2, respectively. The signal was further enhanced in A549-mink ACE2 or A549-ferret ACE2 cells inoculated with SARS-CoV-2pp Y453F (110- or 120-fold, respectively, compared to A549-mouse ACE2). These results suggest that the adaptive Y453F mutation can enhance interaction with *Mustela* ACE2 and consequently promote virus entry.

**Figure 5.**
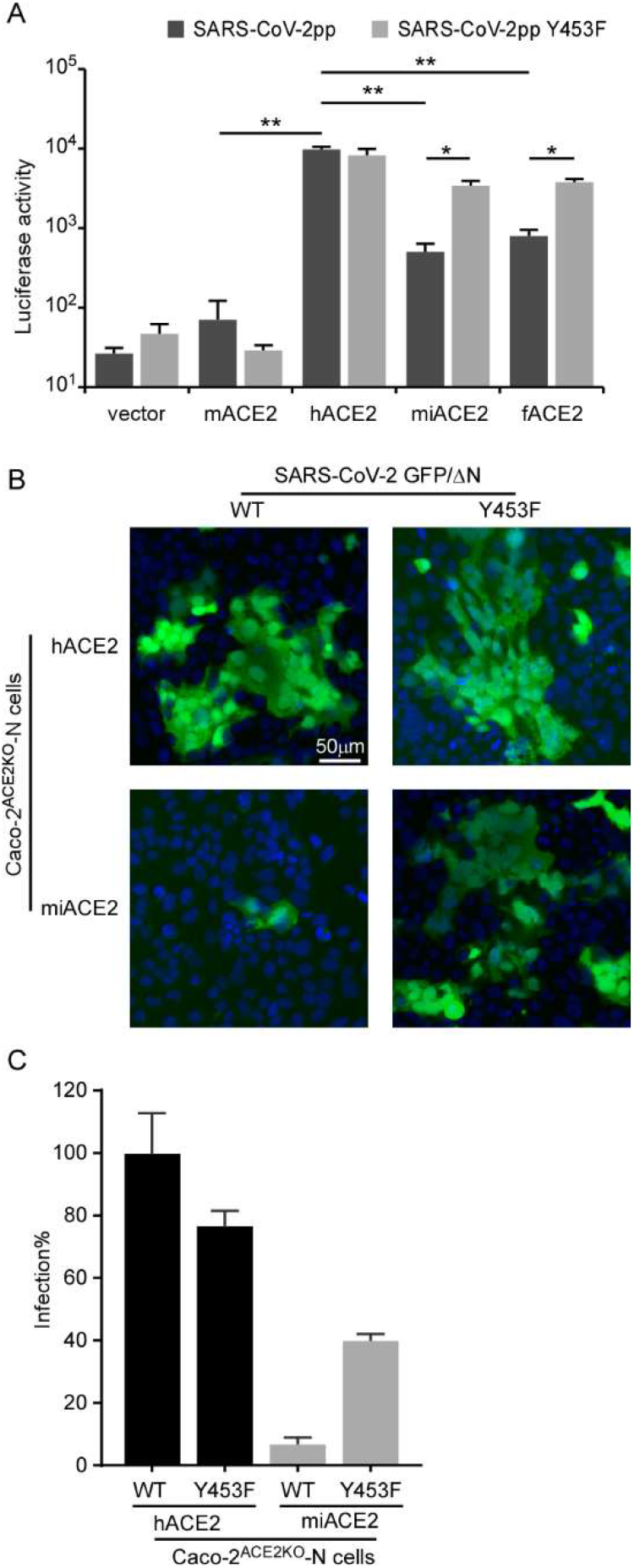
Enhanced infection of Y453F spike pseudotyped virion or trVLP in Caco-2 cells expressing exogenous mink ACE2. (A) HeLa cells transduced with mouse ACE2, human ACE2, mink ACE2 or ferret ACE2 were infected with WT or Y453F SARS-CoV-2 pseudoparticles. Two days after infection, cells were lysed and luciferase activity determined. All infections were performed in triplicate, and the data are representative of two independent experiments (mean ± SD). *, P < 0.05, **, P < 0.01. Significance assessed by one-way ANOVA. (B-C) Endogenous ACE2 knockout Caco-2 cells expressing SARS-CoV-2 N protein were infected with SARS-CoV-2 GFP/ΔN (MOI=0.1). After 36 hours, GFP expression was observed using immunofluorescence microscopy and GFP-positive cells were quantified by flow cytometry. All infections were performed in triplicate, and the data are pooled from two independent experiments (mean ± SD). *, P < 0.05, **, P < 0.01. Significance assessed by one-way ANOVA.

In addition, utilizing the recently developed SARS-CoV-2 transcription and replication-competent virus-like particles (trVLP) cell culture system, in which the SARS-CoV-2 complete life cycle is recapitulated in cells expressing SARS-CoV-2 N gene *in trans*^26^, we produced SARS-CoV-2 GFP/ΔN trVLPs (WT or Y453F). We knocked out endogenous ACE2 in Caco-2 cells expressing the viral N gene (Caco-2-N) using the CRISPR-Cas9 system^27^, and then overexpressed human (Caco-2^ACE2KO^-N-hACE2) or mink ACE2 (Caco-2^ACE2KO^-N-miACE2) (**Fig. S6**). Next, we inoculated the Caco-2 ^ACE2KO^-N-hACE2 and Caco-2 ^ACE2KO^-N-miACE2 cells with the same amount of SARS-CoV-2 GFP/ΔN trVLP or SARS-CoV-2 GFP/ΔN (Y453F) trVLP to achieve similar levels of initial infection. After 36 hours, the cells were collected and virus infection, as evidenced by GFP expression, was quantified by flow cytometry (**Fig. 5B**). As expected, SARS-CoV-2 trVLP replicated and spread well over time in Caco-2 ^ACE2KO^-N-hACE2 cells but not in Caco-2 ^ACE2KO^-N-miACE2 cells. By contrast, SARS-CoV-2 (Y453F) trVLP replicated and spread well in both cell lines (**Fig. 5B**), consistent with our binding experiments and pseudotyped virons assays (**Fig. 2B, 2E, 4A** and **Fig. 5A**). In sum, these results confirm that Y453F is an adaptive mutation in the SARS-CoV-2 spike protein that dramatically enhances utilization of mink ACE2 for entry, conferring a fitness advantage in the mink host.

### Sensitivity of miSARS-CoV-2 variant to neutralization by convalescent sera and ACE2-Ig

SARS-CoV-2 infection elicits neutralizing antibodies that target the spike protein and are critical for protection from re-infection^28^. Given that the mink variant contains several mutations in the spike protein, these mutations might contribute to the evasion of neutralizing antibodies. Therefore, we sought to evaluate the sensitivity of SARS-CoV-2 mink variant to neutralization by convalescent serum. We collected convalescent sera from 10 patients who contracted SARS-CoV-2 (**Fig. 5A**). We produced MLV virons pseudotyped with WT or Y453F spike and then measured the neutralization activity of convalescent sera against these viruses (**Fig. 5B**). The results showed that neutralization of particles bearing the Y453F S protein was almost 3.5-fold less efficient than those bearing the WT protein. This result suggests that the mink variant might evade neutralization by antibodies that mediate protection against COVID-19 in convalescent patients.

Soluble human ACE2 protein can bind to the S protein of SARS-CoV-2 and thus block the interaction between viral spike protein and cell-surface ACE2, thereby decreasing entry of the virus into tissue^29^. As the miSARS-CoV-2 spike exhibited increased binding affinity with human ACE2, it is conceivable that it could be effectively blocked by soluble human ACE2 protein. To this end, we determined the antiviral potency of a recombinant ACE2-Ig variant (human ACE2 aa18-740 fused to an lgG Fc fragment) against WT or mink variant SARS-CoV-2 (Cluster Five) spike (**Fig. 1A**). As shown in **Fig. 6B**, ACE2-Ig effectively inhibited virions pseudotyped with either WT or mink variant spike at an IC_50_ of 62.7ng/ml and 65.7ng/ml, respectively.

**Figure 6.**
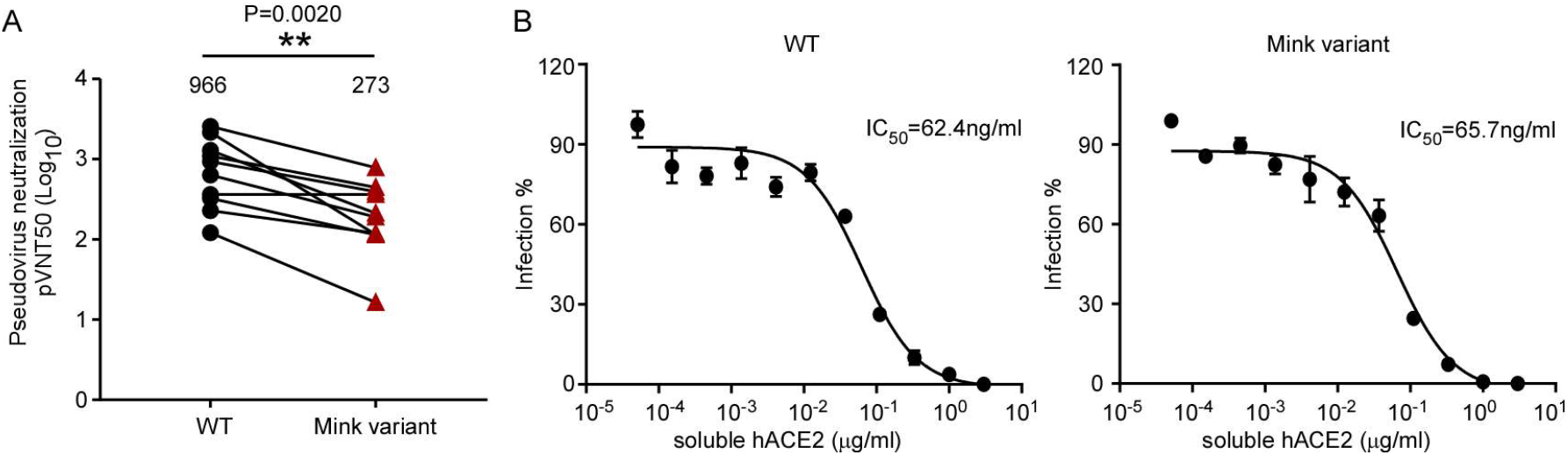
Sensitivity of miSARS-CoV-2 “Cluster Five” spike pseudotyped virus to neutralization by convalescent sera of patients or soluble ACE2. (A) Convalescent serum samples from donors were serially diluted and incubated with WT or “Cluster Five” spike-pseudotyped viruses. The serum dilution factor leading to 50% reduction of pseudotyped virion entry was calculated as the NT_50_ (neutralizing titer 50). Identical plasma samples are connected with lines. These analyses were repeated twice with similar results. Statistical analysis of the difference between neutralization of the WT and “Cluster Five” was performed using two-tailed Wilcoxon matched-pairs signed-rank test. (B) Recombinant ACE2-Ig was diluted at the indicated concentrations. Viral entry was determined by assessing Luc activity 48 hours post infection. The dilution factors leading to 50% reduction of pseudotyped virion entry was calculated as the IC_50_. Data shown are representative of three independent experiments with similar results, and data points represent mean ± SD in triplicate.

Taken together, our data demonstrate that the miSARS-CoV-2 spike potentially evolved to escape neutralization by convalescent serum but still remains sensitive to an ACE2-based therapeutic strategy.

## Discussion

Minks can act as a reservoir of SARS-CoV-2, circulating the virus among them and thus becoming a source for virus spill-over into humans as observed in the Netherlands^14,18^. The miSARS-CoV-2 variants harbor mutations in the spike protein, which potentially could alter its transmission, pathogenicity, and/or sensitivity to vaccine-elicited immunity. In this study, we demonstrated that the Y453F substitution in the miSARS-CoV-2 spike protein is an adaptive mutation that significantly enhances interaction with mink ACE2 and promotes infection of minks. Moreover, the Y453F substitution does not compromise the ability of the spike protein to utilize human ACE2, explaining at least in part the circulation of the mink-associated virus observed in humans. The miSARS-CoV-2 variant exhibited partial resistance to neutralization by convalescent sera, suggesting that the variant has the potential to also escape protection induced by infection or vaccines.

The interaction of spike with ACE2 is a major genetic determinant of SARS-CoV-2 host range^4,7,30,31^, and thus changes in the spike protein can adapt the virus to new hosts. Our findings revealed that SARS-CoV-2 was genetically flexible in its ability to adapt to new hosts. As we have shown, SARS-CoV-2 spike has limited binding capability to mink ACE2 (**Fig. 2B** and **E**), and our previous work demonstrated that mink ACE2 supported SARS-CoV-2 entry at 20% the efficiency of human ACE2^7^. It is conceivable that after transmission from humans to minks, the virus evolved further in mink to adapt to the new host, with efficient binding of SARS-CoV-2 S protein to mink ACE2 a prerequisite for ready transmission among minks. Of interest, a recent study reported the absence of natural SARS-CoV-2 human-to ferret transmission in a high-exposure setting, and genetic analysis suggested that infection of ferrets may require viral adaptation^32^. Here we demonstrate that the emergence of Y453F in SARS-CoV-2 spike significantly enhanced its interaction with other *Mustela* ACE2 orthologs—namely ferret and stoat—conferring a potential fitness advantage in *Mustela* species that could subsequently promote virus transmission and possible risk of animal-adapted virus with genetic alterations to spilling over into humans.

Since the spike protein is a major target for prophylactic vaccines and antibody-based therapeutics, such mutations could have implications for treatment, diagnostic tests and viral antigenicity. It has been shown that the D614G substitution in the SARS-CoV-2 spike can promote virus entry into host cells and enhance infectivity as well as make the mutant virus resistant to neutralizing antibody^33,34^. The Y453F mutation was shown to be an escape mutation for the monoclonal antibody RGN10933^35^. Our results showed that miSARS-CoV-2 Y453F spike pseudotyped virions exhibited 3.5-fold resistance to neutralization by convalescent serum (**Fig. 6A**), suggesting the antibody response induced by infection or vaccine might not offer sufficient protection against mink-associated SARS-CoV-2 variant infection. However, the miSARS-CoV-2 was still sensitive to ACE2-lg, which suggests that ACE2-based therapeutics may represent an effective antiviral strategy against the continued emergence of new variants.

In summary, our study suggests that Y453F is an adaptive mutation in SARS-CoV-2 that results in a virus more competent for infection and transmission among mink. In addition, the miSARS-CoV-2 Y453F spike mutant maintained its ability to interact with human ACE2 and exhibited partial resistance to neutralizing antibodies, potentially explaining the ability of miSARS-CoV-2 to transmit back to humans and subsequent human-human transmission. Thus, it remains an urgent need to identify potential zoonotic reservoirs of SARS-CoV-2 and to monitor the evolution of the virus in zoonotic hosts to prevent future outbreaks and develop appropriate countermeasures.

## ACKNOWLEDGEMENTS

We wish to acknowledge Di Qu, Zhiping Sun, Wendong Han and other colleagues at the Biosafety Level 3 Laboratory of Fudan University for help with experiment design and technical assistance. We thank the staffs from BL17B/BL18U1/BL19U1/BL19U2/BL01B beamline of National Facility for Protein Science in Shanghai (NFPS) at Shanghai Synchrotron Radiation Facility, for assistance during data collection. We thank Dr. Jenna M. Gaska for suggestions and revision of the manuscript.

This work was supported by National Natural Science Foundation of China (32041005), Tsinghua-Peking University Center of Life Sciences (045-61020100120), Start-up Foundation of Tsinghua University (53332101319).

## MATERIALS AND METHODS

### Cell culture

HEK293T (American Tissue Culture Collection, ATCC, Manassas, VA, CRL-3216), A549 (ATCC #CCL-185), Vero E6 (Cell Bank of the Chinese Academy of Sciences, Shanghai, China) and HeLa (ATCC #CCL-2) cells were maintained in Dulbecco’s modified Eagle medium (DMEM) (Gibco, NY, USA) supplemented with 10% (vol/vol) fetal bovine serum (FBS), 10mM HEPES, 1mM sodium pyruvate, 1×non-essential amino acids, and 50 IU/ml penicillin/streptomycin in a humidified 5% (vol/vol) CO_2_ incubator at 37°C. Cells were tested routinely and found to be free of mycoplasma contamination.

### Plasmids

The cDNAs encoding human or mink ACE2 were synthesized by GenScript and cloned into pLVX-IRES-zsGreen1 vectors (Catalog No. 632187, Clontech Laboratories, Inc) with a C-terminal FLAG tag. ACE2 mutants were generated by Quikchange (Stratagene) site-directed mutagenesis. All of the constructs were verified by Sanger sequencing.

### Lentivirus production

Vesicular stomatitis virus G protein (VSV-G) pseudotyped lentiviruses expressing ACE2 orthologs tagged with FLAG at the C-terminus were produced by transient co-transfection of the third-generation packaging plasmids pMD2G (Addgene #12259) and psPAX2 (Addgene #12260) and the transfer vector with VigoFect DNA transfection reagent (Vigorous) into HEK293T cells. The medium was changed 12 h post transfection. Supernatants were collected at 24 and 48h after transfection, pooled, passed through a 0.45-μm filter, and frozen at −80°C.

### Western blotting

Sodium dodecyl sulfate-polyacrylamide gel electrophoresis (SDS-PAGE) immunoblotting was performed as follows: After trypsinization and cell pelleting at 2,000 × g for 10 min, whole-cell lysates were harvested in RIPA lysis buffer (50 mM Tris-HCl [pH 8.0], 150mM NaCl, 1% NP-40, 0.5% sodium deoxycholate, and 0.1% SDS) supplemented with protease inhibitor cocktail (Sigma). Lysates were electrophoresed in 12% polyacrylamide gels and transferred onto nitrocellulose membrane. The blots were blocked at room temperature for 0.5 h using 5% nonfat milk in 1× phosphate-buffered saline (PBS) containing 0.1% (v/v) Tween 20. The blots were exposed to primary antibodies anti-β-Tubulin (CW0098, CWBIO), or anti-FLAG (F7425, Sigma) in 5% nonfat milk in 1× PBS containing 0.1% Tween 20 for 2 h. The blots were then washed in 1× PBS containing 0.1% Tween 20. After 1h exposure to HRP-conjugated secondary antibodies, subsequent washes were performed and membranes were visualized using the Luminescent image analyzer (GE).

### Surface ACE2 binding assay

HeLa cells were transduced with lentiviruses expressing the ACE2 from different species for 48 h. The cells were collected with TrypLE (Thermo #12605010) and washed twice with cold PBS. Live cells were incubated with the recombinant protein, S1 domain of SARS-CoV-2 spike C-terminally fused with Fc (Sino Biological #40591-V02H, 1μg/ml) at 4°C for 30 min. After washing, cells were stained with goat anti-human IgG (H + L) conjugated with Alexa Fluor 647 (Thermo #A21445, 2 μg/ml) for 30 min at 4°C. Cells were then washed twice and subjected to flow cytometry analysis (Thermo, Attune™ NxT).

### Production of SARS-CoV-2 pseudotyped virus

Pseudoviruses were produced in HEK293T cells by co-transfecting the retroviral vector pTG-MLV-Fluc, pTG-MLV-Gag-pol, and pcDNA3.1 expressing SARS-CoV-2 spike gene or VSV-G (pMD2.G, Addgene #12259) using VigoFect (Vigorous Biotechnology). At 48 h post transfection, the cell culture medium was collected for centrifugation at 3500 rpm for 10 min, and then the supernatant was subsequently aliquoted and stored at −80°C for further use. Virus entry was assessed by transduction of pseudoviruses in A549 cells expressing ACE2 orthologs or mutants in 48-well plates. After 48 h, intracellular luciferase activity was determined using the Luciferase Assay System (Promega, Cat. #E1500) according to the manufacturer’s instructions. Luminescence was recorded on a GloMax® Discover System (Promega).

### Protein expression and purification

The SARS-CoV-2 RBD (residues Arg319–Phe541) and the N-terminal peptidase domain of human ACE2 (residues Ser19–Asp615) were expressed using the Bac-to-Bac baculovirus system (Invitrogen) as described previously^21^. ACE2-lg, a recombinant Fc fusion protein of soluble human ACE2 (residues Gln18-Ser740), was expressed in 293F cells and purified using protein A affinity chromatography as described in our previous study^36^.

### Surface plasmon resonance anlysis

ACE2 was immobilized on a CM5 chip (GE Healthcare) to a level of around 500 response units using a Biacore T200 (GE Healthcare) and a running buffer (10 mM HEPES pH 7.2, 150 mM NaCl and 0.05% Tween-20). Serial dilutions of the SARS-CoV-2 RBD were flowed through with a concentration ranging from 400 to 12.5 nM. The resulting data were fit to a 1:1 binding model using Biacore Evaluation Software (GE Healthcare).

### Protein crystallization

The gene encoding 333-527aa of SARS-CoV-2 RBD (WT or Y453F mutant) was cloned into the pFastbac-dual vector using the BamH1 and Hind3 restriction enzyme. A GP67 signal peptide was added to the N-terminus for protein secretion and a 6×His tag was added to the C-terminus for protein purification as described previously^21^. The gene encoding 1-615aa of ACE2 from human or mink was constructed in the same way as SARS-CoV-2 RBD except using its own peptide for protein secretion. All the proteins were expressed using Hi5 cells. Briefly, baculovirus containing the gene of SARS-CoV-2 RBD or ACE2 were produced according to the Invitrogen bac-to-bac Baculovirus Expression System Manual. To purify the protein, supernatant was collected 60h after cell transfection, then buffer-exchanged to 1×HBS (10mM HEPES, 150mM NaCl, pH7.2), and was loaded on a Ni column equilibrated with the same buffer. After washing the column with 1×HBS containing 20mM Imidazole, the protein was eluted by 1×HBS containing 500mM Imidazole. The protein was further concentrated and purified by gel filtration chromatography on a Superdex 200 10/300 column in the 1 ×HBS buffer. To prepare the RBD and ACE2 complex, ACE2 and RBD were mixed as a molar ratio of 1:1.5 and incubated at 4°C for 60 minutes, then purified by gel filtration chromatography in TBS buffer (25mM Tris, 150mM NaCl, pH 7.5). Fractions were collected and concentrated to 13mg/ml for crystal screening. Kits used in crystal screening were all purchased from Hampton Research. The crystal of SARS-CoV-2 RBD Y453F and mink ACE2 complex was obtained in a well solution of 1.6 M sodium/potassium phosphate using the sitting drop method.

### Data collection and refinement

Diffraction data were collected at 100 K and at a wavelength of 0.9796 Å on the BL18U1 beam line of the Shanghai Synchrotron Research Facility. The diffraction data were then processed using the HKL2000. The two structures were determined using the molecular replacement method with Molrep in the CCP4 suite^37^, using the model of the human ACE2 and SARS-CoV-2 complex^21^. There were three complex molecules in one asymmetric unit. Subsequent model building and refinement were performed using COOT and PHENIX, respectively^38,39^. The data-processing statistics and structure refinement statistics are listed in **Table 1**. All structure figures were generated with PyMol^40^. The coordinates and structure factor files for the SARS-CoV-2 Y453F RBD/mink ACE2 complex have been deposited in the Protein Data Bank (PDB) under accession number 7F5R.

### Convalescent serum

We obtained convalescent serum from 10 patients (**Table S2**, one patient’s information is not complete) more than one month after documented SARS-CoV-2 infection in the spring of 2020. Among these 10 patients, 9 patients information was recorded. Three had severe disease and 6 had non-severe disease. Their ages ranged from 24 to 69, with a mean of 54. Eight were male, and 1 was female. Each plasma sample was heat-inactivated (56°C, 30 min) and then assayed for neutralization against WT or mink-variant pseudoviruses. This study was approved by the Institution Review Board of Tsinghua University (20210040).

### SARS-CoV-2 pseudovirus neutralization assay

SARS-CoV-2 spike variant-pseudotyped luciferase reporter viruses were pre-diluted in DMEM (2% FBS, heat-inactivated) containing serially diluted convalescent serum. Virus-serum mixtures were incubated at 37°C for 30min, then added to A549-hACE2 cells in 96-well plates and incubated at 37°C. After 48 h, intracellular luciferase activity was determined using the Luciferase Assay System (Promega, Cat. #E1500) according to the manufacturer’s instructions. Luminescence was recorded on a GloMax Discover System (Promega). The NT50 (Neutralizing titer 50) was defined as the serum dilution factor that leads to 50% reduction in spike protein-driven cell entry.

### Statistics analysis

One-way analysis of variance (ANOVA) with Tukey’s honestly significant difference (HSD) test was used to test for statistical significance of the differences between the different group parameters. *P* values of less than 0.05 were considered statistically significant.

### SUPPLEMENTAL FIGURES AND FIGURE LEGENDS

**Supplemental Figure 1.**
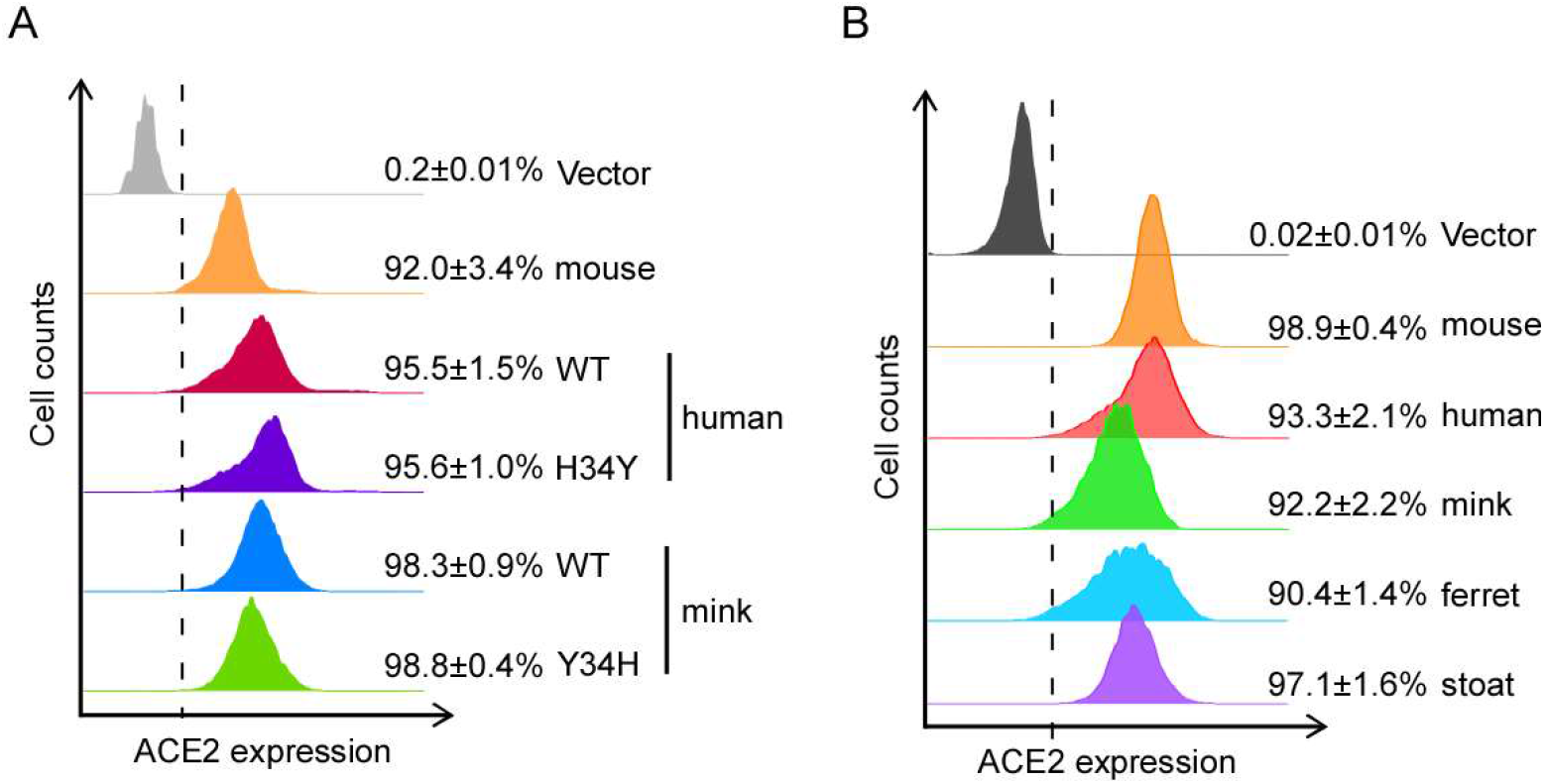
Cell surface expression of human and mink ACE2. (A) HeLa cells transduced with lentiviruses (pLVX-IRES-zsGreen) expressing mouse, human (WT or H34Y) or mink ACE2 (WT or Y34H) were incubated with rabbit polyclonal antibody (Sino Biological Inc. China, Cat: 10108-T24) against ACE2. The cells were washed and then stained with 2μg/mL goat anti-rabbit IgG (H+L) conjugated with Alexa Fluor 568 for flow cytometry analysis. The cell surface ACE2 was calculated as the percent of Alex Fluor 568-positive cells among the zsGreen-positive cells. This experiment was repeated twice with similar results. (B) HeLa cells transduced with lentiviruses (pLVX-IRES-zsGreen) expressing mouse, human, mink, ferret or stoat ACE2 were analyzed as described in (A) panel. This experiment was repeated twice with similar results.

**Supplemental Figure 2.**
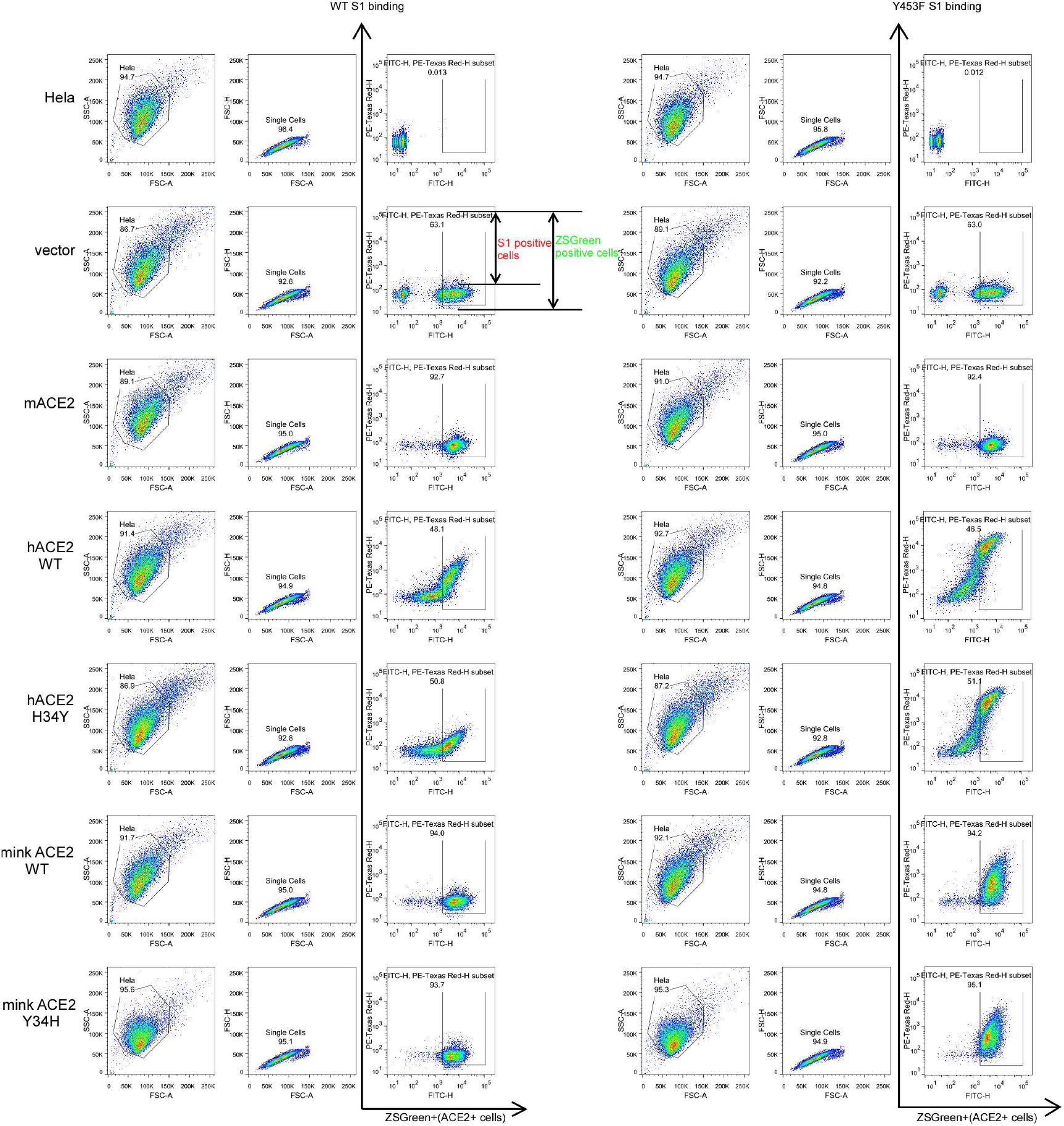
Gating strategy for determining the binding efficiency of different ACE2 orthologs with SARS-CoV-2 S1 protein. The main cell population was identified and gated on Forward and Side Scatter. Single cells were further gated on FSC-A and FSC-H. The gated cells were plotted by FITC-H (zsGreen, the ACE2-expressing population) and APC-H or PE-Texas Red-H (S1-Fc bound population). The APC-H or PE-Texas Red-H positive cell population was plotted as a histogram to show the S1-Fc positive population as in Fig 2A. The binding efficiency was defined as the percent of S1-Fc binding cells among the zsGreen-positive cells. Shown are FACS plots representative of those used for the calculations of binding efficiencies of ACE2 orthologs with S1-Fc. All binding assays were performed in duplicate.

**Supplemental Figure 3.**
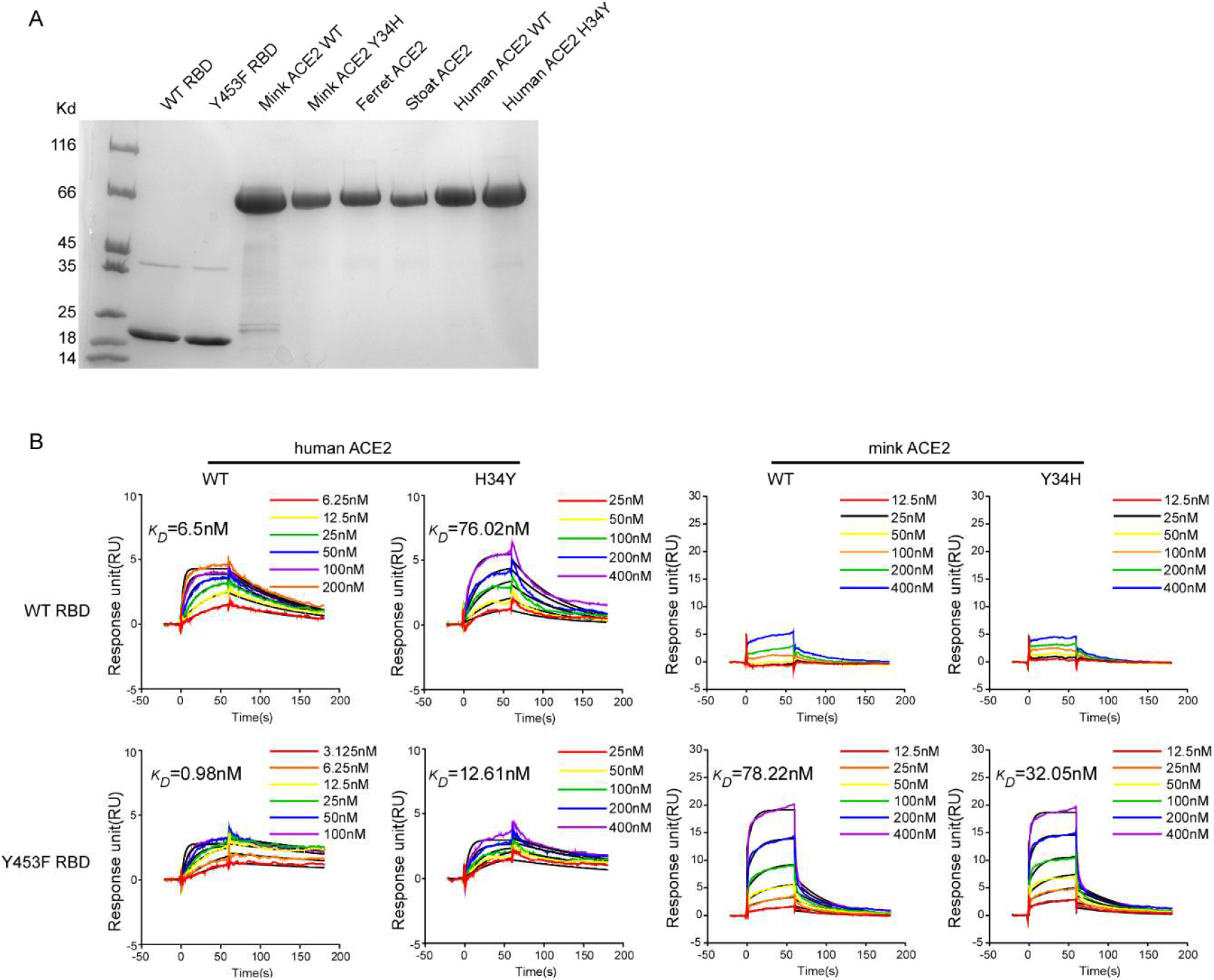
SPR analysis of the binding affinity of ACE2 variants with WT or Y453F RBD. (A) The N-terminal peptidase domain of each ACE2 variant (residues Met1-Asp615), WT or Y453F SARS-CoV-2 RBD (residues Thr333-Pro527) were expressed and purified as described in the *Materials and Methods*. The purified proteins were analyzed by SDS-PAGE with Coomassie blue staining. (B) The binding kinetics of ACE2 variant proteins (human or mink) with recombinant WT or Y453F SARS-CoV-2 RBD were obtained using the BIAcore. ACE2 proteins were captured on the chip, and serial dilutions of RBD were then injected over the chip surface. Experiments were performed three times with similar results, and one set of representative data is displayed.

**Supplemental Figure 4.**
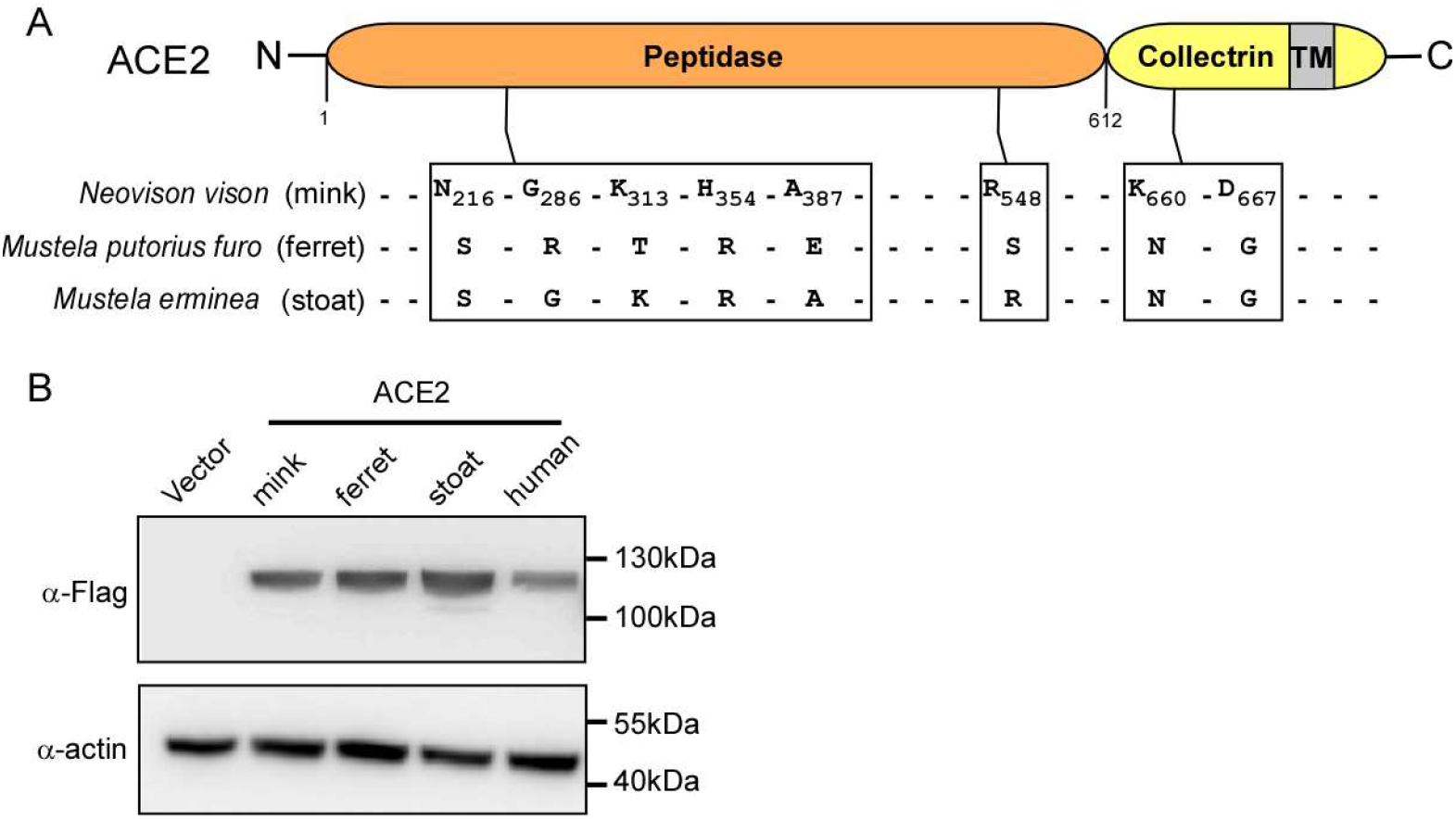
Expression of *Mustelidae* species ACE2 orthologs in HeLa cells. (A) Alignment of ACE2 orthologs of the *Mustelidae* species mink, ferret and stoat. (B) Representative immunoblots of HeLa cells transduced with lentiviruses expressing FLAG-tagged ACE2 orthologs as indicated. Actin used as the loading control. These experiments were independently performed twice with similar results.

**Supplemental Figure 5.**
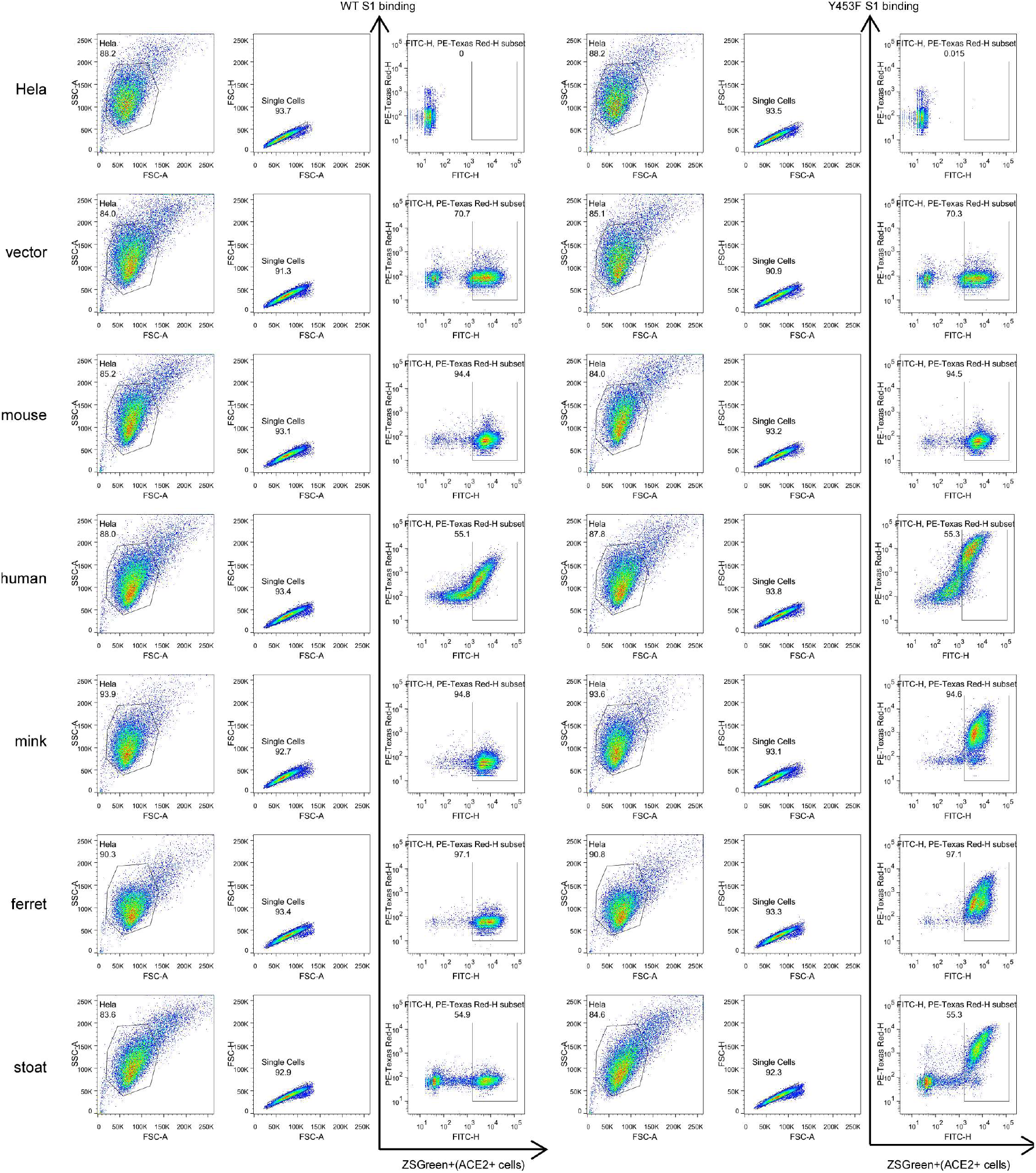
Gating strategy for determination of the binding efficiency of *Mustelidae* species ACE2 with SARS-CoV-2 S1 protein. The main cell population was identified and gated on Forward and Side Scatter. Single cells were further gated on FSC-A and FSC-H. The gated cells were plotted by FITC-H (zsGreen, as the ACE2-expressing population) and APC-H or PE-Texas Red-H (S1-Fc bound population). The APC-H or PE-Texas Red-H positive cell population was plotted as a histogram to show the S1-Fc positive population as in Fig 2D. The binding efficiency was defined as the percent of S1-Fc binding cells among the zsGreen-positive cells. Shown are FACS plots representative of those used for the calculations of binding efficiencies of ACE2 orthologs with S1-Fc.

**Supplemental Figure 6.**
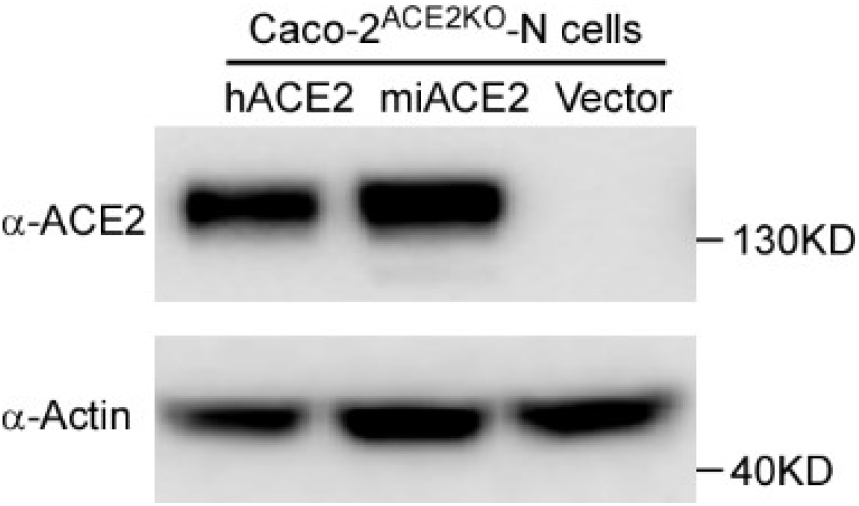
Ectopic expression of human ACE2 and mink ACE2 in Caco-2^ACE2KO^-N cells. Caco-2^ACE2KO^-N cells were transduced with lentivirus expressing human ACE2 or mink ACE2. Immunoblotting assay was performed to detect the expression of human ACE2 and mink ACE2. Actin was used as the loading control. These experiments were independently performed twice with similar results.

### SUPPLEMENTAL TABLES

**Table S1.** Comparison of contact residues on the interfaces of SARS-CoV-2 WT RBD/human ACE2 and Y453F RBD/mink ACE2.

| Y453F RBD | Mink ACE2 | WT RBD | Human ACE2 |
| --- | --- | --- | --- |
| **R403** | Y34 | **R403** |  |
| **K417** | E30 | **K417** | D30 |
| **G446** |  | **G446** | Q42 |
| **Y449** | E38, Q42 | **Y449** | D38, Q42 |
| **F453** | Y34 | **Y453** | H34 |
| **L455** | Y34 | **L455** | H34 |
| **F456** | T27, E30 | **F456** | T27, D30 |
| **A475** | S19, L24 | **A475** | Q24, T27 |
| **G476** | L24 | **G476** |  |
| **F486** |  | **F486** | L79, M82, Y83 |
| **N487** | L24, Y83 | **N487** | Q24, Y83 |
| **Y489** | T27, F28, Y83 | **Y489** | T27, F28, K31, Y83 |
| **Q493** | K31, Y34 | **Q493** | K31, H34, E35 |
| **Y495** | E38 | **Y495** |  |
| **G496** | E38, K353 | **G496** | K353 |
| **Q498** | E38, Y41, Q42, K353 | **Q498** | Y41, Q42 |
| **T500** | Y41, N330, D355, R357 | **T500** | Y41, N330, D355, R357 |
| **N501** | Y41, K353 | **N501** | Y41, K353 |
| **G502** | K353, H354, D355 | **G502** | K353, G354 |
| **Y505** | E37, K353, H354, R393 | **Y505** | E37, K353, G354, R393 |

A distance of 4 Å was defined as interaction residues.

**Table S2.** Summary of COVID-19 patients’ information.

| **Characteristics** | **Patients (n=9)** |
| --- | --- |
| **Age (median, range)** | 54 (24-69) |
| **Sex** |  |
| Male (%) | 8 (88.9) |
| Female (%) | 1 (11.1) |
| **Disease severity** |  |
| Severe | 3 (33.3) |
| Non-severe | 6 (66.7) |
| **Comorbidities, n (%)** |  |
| Fever | 6 (66.7) |
| Fatigue | 4 (44.4) |
| Dry cough | 2 (22.2) |
| Dyspnea | 3 (33.3) |
| Expectoration | 4 (44.4) |
| Diarrhea | 1 (11.1) |
| Chills | 2 (22.2) |
| Chest stuffiness | 3 (33.3) |

